# Discovery of MA48, a Small Molecule Inhibitor of CAPON (NOS1AP)-NOS1 Protein-Protein Interaction

**DOI:** 10.64898/2026.02.21.707155

**Authors:** Ashraf N. Abdo, Hossam Nada, Sungwoo Cho, Moustafa Gabr

## Abstract

CAPON (also known as NOS1AP) is an adaptor protein of neuronal nitric oxide synthase (nNOS) that has been implicated in the progression of multiple neurodegenerative diseases, making it an attractive but largely unexplored therapeutic target. To identify small molecule CAPON modulators, we screened a library of 10,000 compounds for CAPON binding using affinity selection-mass spectrometry (AS-MS), which led to the identification of compound **MA48** as a potential CAPON binder. Subsequent biophysical validation using microscale thermophoresis (MST) confirmed direct binding, with **MA48** exhibiting a dissociation constant (Kd) of 11.9 µM. Structure–activity relationship (SAR) analysis combined with molecular docking was performed to elucidate key pharmacophoric features underlying the **MA48**/CAPON interaction. To determine whether **MA48** disrupts the CAPON-nNOS interaction in a cellular context, we conducted a NanoBRET assay, which demonstrated that **MA48** significantly inhibited this interaction in living cells. Collectively, these findings suggest that **MA48** represents the first reported small molecule inhibitor of CAPON and provides a foundation for further development of CAPON-targeted therapeutics.

## 1. Introduction

CAPON (also known as NOS1AP) is a neuronal adaptor protein that interacts with neuronal nitric oxide synthase (nNOS), playing a central role in the regulation of nitric oxide (NO) signaling and diverse physiological processes within the nervous system. Through its C-terminal PDZ-binding motif, CAPON binds to the N-terminal PDZ domain of nNOS, modulating NO production and downstream signaling pathways involved in synaptic plasticity, apoptosis, and neuronal development [1-3]. Under normal conditions, nNOS associates with PSD-95 to form a signaling complex linked to NMDA receptors, where glutamate-induced calcium influx activates nNOS and stimulates NO generation essential for learning and memory[1, 4, 5]. However, excessive glutamatergic activation can overdrive this system, producing neurotoxic levels of NO and reactive nitrogen species[6, 7] (Figure 1).

**Figure 1.**
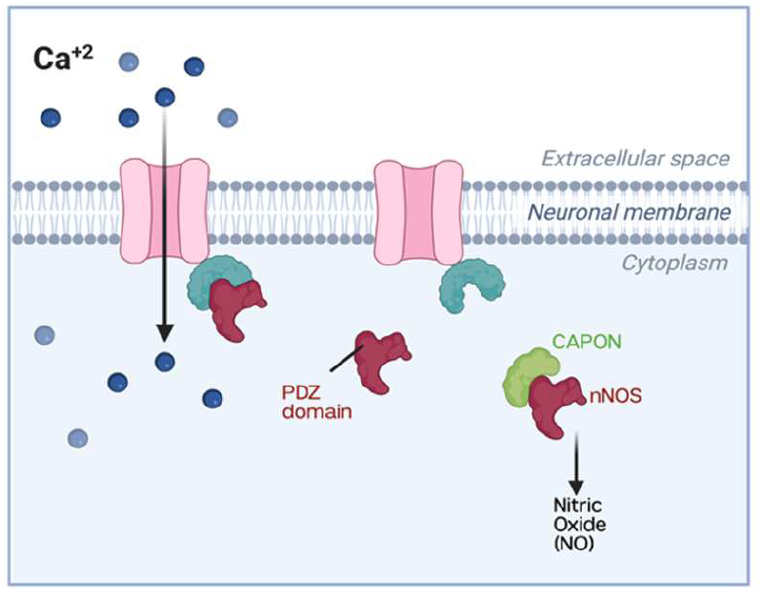
Illustrating the functional interaction between neuronal nitric oxide synthase (nNOS), its PDZ domain, and CAPON at the neuronal membrane. Calcium ions (Ca^2+^) enter the neuron through a membrane channel, triggering intracellular signaling events. The PDZ domain of nNOS is depicted as the primary interaction interface that associates with CAPON in the cytoplasm. CAPON binds to nNOS and modulates its activity, ultimately influencing the production of nitric oxide (NO). Emphasizing how CAPON overexpression be linked to abnormal nitric oxide signaling through the nNOS-CAPON complex.

CAPON can compete with PSD-95 for binding to nNOS, forming an alternative complex that reduces NO synthesis and protects neurons from excitotoxic damage[8]. While this modulation is beneficial under stress conditions, abnormal overexpression of CAPON disrupts physiological nNOS–PSD-95 signaling and impairs normal neuronal communication[1]. Dysregulation of this regulatory axis has been implicated in numerous neurological and psychiatric disorders, including Alzheimer’s, Parkinson’s, and Huntington’s diseases, amyotrophic lateral sclerosis, depression, bipolar disorder, and anxiety-related conditions [9-14]. Mechanistically, elevated CAPON levels have been linked to pathological Tau phosphorylation, increased β-amyloid deposition, and formation of the Dexras1–nNOS–CAPON complex molecular events that drive neuronal loss and synaptic dysfunction [8, 15]. Altered CAPON activity also perturbs MAPK signaling, leading to decreased ERK and increased p38 activation, further contributing to neurodegenerative outcomes [16].

Given its involvement in these critical disease pathways, CAPON has emerged as a compelling and underexplored therapeutic target. Despite its well-established role in neuronal signaling and neurodegeneration, CAPON has remained largely unexplored from a small molecule discovery perspective. Unlike enzymes with well-defined catalytic pockets, CAPON functions primarily as an adaptor protein mediating protein-protein interactions, which are traditionally considered challenging targets due to extended, shallow binding interfaces and limited structural information. The absence of experimentally resolved high-resolution structures further complicates rational ligand design, contributing to the perception that CAPON may be chemically intractable.

While genetic and peptide-based strategies have provided insights into CAPON-dependent signaling, small molecules offer distinct advantages, including cell permeability, reversible modulation, tunable pharmacokinetics, and compatibility with in vivo applications. To date, no validated small molecule CAPON inhibitors have been reported, and the chemical tractability of the CAPON-NOS1 interface remains unknown. Establishing whether CAPON can be engaged by drug-like small molecules is therefore a critical step toward therapeutic development.

To explore whether CAPON can be modulated by small molecules, we applied an affinity selection–mass spectrometry (AS-MS)-based screening strategy, a label-free platform that enables direct detection of protein-ligand interactions in solution. Given the limited structural information available for CAPON, the AS-MS screening strategy is capable of detecting direct protein-ligand interactions in solution without prior knowledge of binding pockets. This methodology is particularly well-suited for challenging adaptor proteins where conventional high-throughput functional assays may not be readily available. This approach identified a series of CAPON-binding candidates from a diverse compound library, which were further validated using microscale thermophoresis (MST) to quantify binding affinity. Among them, compound **MA48** displayed measurable binding to CAPON in the low micromolar range, establishing it as confirmed small molecule ligand of CAPON. These findings provide the first evidence that CAPON can be directly engaged and functionally modulated by small molecules, establishing a foundation for structure-guided optimization and therapeutic exploration of CAPON-dependent signaling pathways.

## 2. Methods and materials

### 2.1 Affinity Selection Mass Spectrometry (AS-MS)

AS-MS Screening was performed as described in the Supporting Information file and previously described [17-19].

### 2.2. Dose-response characterization using Microscale thermophoresis (MST)

Compounds that produced a clear binding response in the single-point screen were subsequently evaluated in a concentration-dependent MST assay. Twelve-point serial dilutions were prepared, spanning a starting concentration of 250 µM down to the low-nanomolar range, and mixed with labeled CAPON-His protein while maintaining a final DMSO concentration of 2.5%. After a 30-minute equilibration step at room temperature, samples were loaded into standard MST capillaries and measured on a Monolith NT.115 instrument, using medium-to-high infrared laser power and 60–80% LED excitation in the red detection channel. Dissociation constants (Kd) were calculated using the MO.Affinity Analysis package (NanoTemper Technologies).

### 2.4 Homology Modeling of CAPON Structure

The three-dimensional structure of CAPON was predicted using homology modeling through the SWISS-MODEL[20] workspace accessed on 02-02-2026. The highest-quality template selected for model building was the PTB domain-containing engulfment adapter protein 1 (PDB ID: 6itu.1.A[21]), which exhibited 29.45% sequence identity with the target sequence, covering 29% of the sequence with a sequence similarity of 0.36. The structure was determined by X-ray diffraction at 2.17 Å resolution. The quality of the generated homology model was assessed using the QMEAN scoring function. The final model achieved a Global Model Quality Estimation (GMQE) score of 0.17 and a QMEAND score of 0.57 ± 0.07, indicating moderate confidence in the predicted structure.

### 2.5 Binding Site Prediction

Potential ligand-binding sites on the CAPON homology model were predicted using the PrankWeb [22] server (https://prankweb.cz/). PrankWeb employs a machine learning-based algorithm to identify and rank potential binding pockets based on structural features, evolutionary conservation, and physicochemical properties. The server analyzes the 3D structure and provides predictions of druggable binding sites with associated confidence scores.

### 2.6 Molecular Docking Studies

Molecular docking of the small molecule **MA48** into the predicted binding site of the CAPON homology model was performed using the induced fit protocol of Schrödinger Suite. The CAPON model was prepared using the Protein Preparation Wizard, which includes adding hydrogen atoms, assigning bond orders, creating disulfide bonds, optimizing hydrogen bond networks, and minimizing the structure using the OPLS4 force field. The final docking poses were visualized and analyzed using Discovery Studio Visualizer (BIOVIA, Dassault Systèmes).

### 2.3 NanoBRET assay for NOS1-CAPON interaction

A NanoBRET-based assay system was developed to monitor NOS1-NOS1AP protein-protein interaction in living cells. Full-length human NOS1 was fused with NanoLuciferase (NanoLuc) at the N-terminus, and full-length human CAPON (NOS1AP) was fused with Venus fluorescent protein at the C-terminus. Both constructs were co-transfected into CHO-K1 cells using a suitable transfection reagent. Following transfection, cells were seeded into white 96-well plates and maintained until approximately 80% confluency was achieved. For compound treatment, growth media was aspirated, cells were rinsed twice with PBS, and HBSS solution was added. Test compounds or vehicle control (DMSO) were then applied at the indicated concentrations for 3 hours. NanoBRET signals were detected by adding NanoLuc substrate, and donor (460 nm) and acceptor (535 nm) emissions were recorded using a Tecan plate reader. The NanoBRET ratio was determined by calculating the acceptor/donor emission ratio and normalizing to the vehicle control.

## 3. Results and Discussion

To identify novel small molecule inhibitors of CAPON, we conducted an affinity selection–mass spectrometry (AS-MS) workflow based on previously established protocols [17-19]. Approximately 10,000 compounds from the ThermoScientific HitFinder library were evaluated. This collection represents a chemically diverse subset of the larger Maybridge Screening Collection and was designed using clustering of Daylight molecular fingerprints and a Tanimoto similarity cutoff of 0.7 to ensure broad coverage of drug-like chemical space. All compounds in the library comply with Lipinski’s criteria for drug-likeness (ClogP ≤ 5, ≤10 hydrogen bond acceptors, ≤5 hydrogen bond donors, and molecular weight ≤ 500 Da) and possess a minimum purity of 90%. The HitFinder collection was selected as an initial screening set due to its high chemical diversity, pre-clustered design, and compliance with drug-like physicochemical parameters. For an adaptor protein target such as CAPON, which lacks a defined catalytic pocket or established ligand scaffolds, broad and unbiased chemical coverage is essential to assess tractability. This diversity-oriented library was therefore well suited for identifying early chemical starting points for CAPON engagement. The use of a chemically diverse yet tractable 10,000-member set also enabled efficient AS-MS screening while maintaining manageable analytical complexity and minimizing redundant chemical space sampling.

In the AS-MS assay, CAPON protein was incubated with pooled small molecule mixtures to allow potential protein-ligand complexes to form. These complexes were separated from free compounds via size-exclusion chromatography (SEC). Bound ligands were then dissociated from the protein, separated by high-performance liquid chromatography (HPLC), and identified using time-of-flight mass spectrometry. Compounds were screened in pooled mixtures to increase throughput while maintaining manageable spectral complexity. Pool composition was designed to minimize mass overlap and ion suppression effects. Parallel control samples lacking CAPON protein were processed under identical conditions to identify nonspecific binders and chromatographic artifacts. Enrichment profiles were reproducible across independent pooled screening runs, supporting the robustness of the AS-MS workflow. Following manual inspection of the identified 124 compounds as potential CAPON binders (1.24% apparent hit rate) to eliminate reactive, unstable, or poorly characterized molecules, single dose screen was (100 μM, n=3) performed using dianthus for the top 50 compounds and compound **MA48** (the most potent CAPON binder, Figure 2A) was selected for further validation. The top 50 candidates were prioritized based on enrichment magnitude and signal reproducibility and were subjected to single-concentration orthogonal validation using MST. Compounds demonstrating the highest enrichment ratios (≥3-fold enrichment over control) and favorable physicochemical properties were prioritized for follow-up, while lower-intensity or structurally redundant candidates were deprioritized at this stage. **MA48** was selected for further study based not only on enrichment intensity but also on its synthetic accessibility and amenability to systematic SAR exploration.

**Figure 2.**
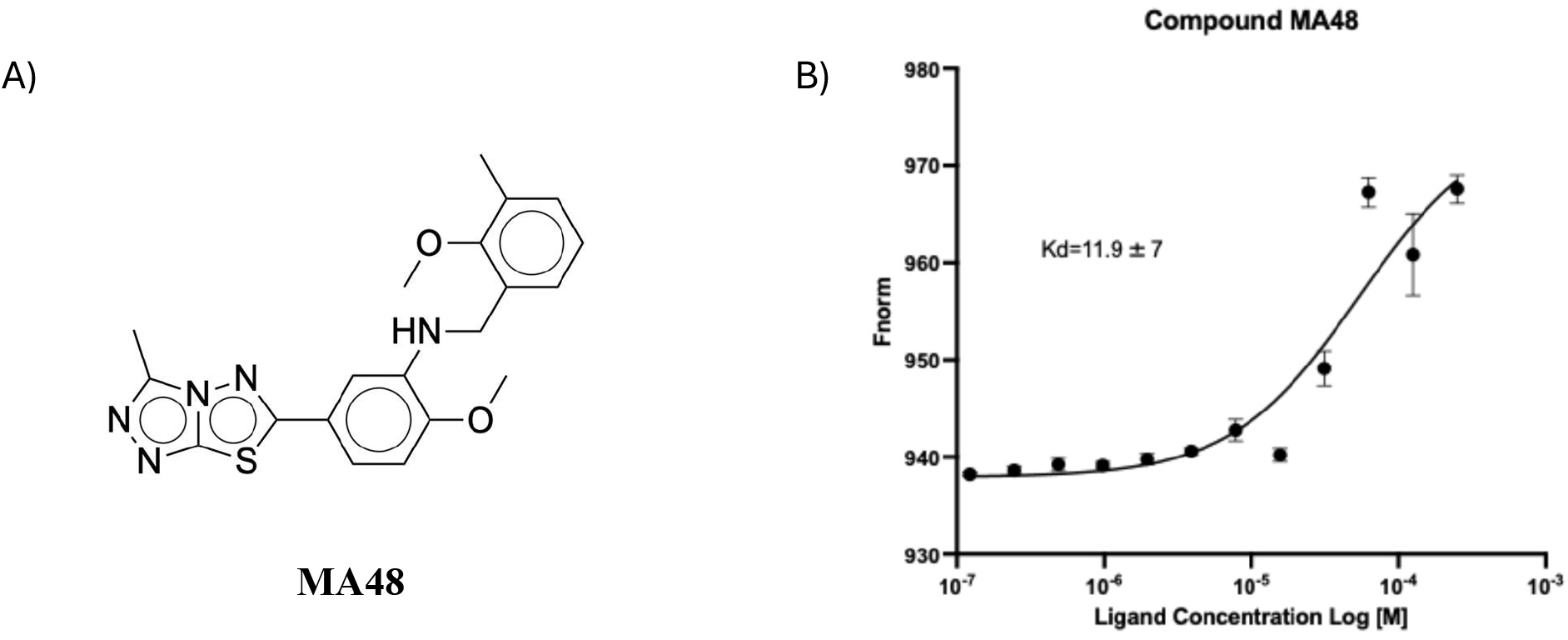
**(A)** Chemical structure of **MA48**, the lead small molecule CAPON inhibitor identified from the AS-MS screen. The compound contains a substituted benzylamine linked to a heterocyclic thiadiazole moiety. **(B)** Dose Response curve of Compound **MA48** with CAPON protein using MST. The graph displays dose-dependent changes in fluorescence (Fnorm) 650 nm plotted against increasing concentration of the compound (n=3, Error bars display SEM).

For MST validation of CAPON binding, His-tagged CAPON labeled with a fluorescent dye was incubated with serial dilutions of each compound prepared in PBST containing 2.5% DMSO. Changes in thermophoretic movement were fitted to the Hill model to derive dissociation constants (Kd). **MA48** displayed a clear dose-dependent binding profile, yielding a Kd of 11.9 ± 7 µM (Figure 2B). This level of affinity is consistent with that typically seen for fragment-sized ligands identified through AS-MS screening strategies. Taken together, these findings demonstrate that **MA48** engages CAPON directly in solution and justify its selection for subsequent structural and mechanistic studies.

### Preliminary structure-activity relationship (SAR) analysis

To assess the chemical tractability of the MA48 scaffold and to validate key structural determinants of CAPON binding, a focused SAR-by-catalog approach was undertaken. A small, structurally related subset of commercially available analogs was selected to probe the contribution of the heterocyclic core, linker functionality, and peripheral substituents to binding affinity. Given that this represents the first reported small molecule chemotype targeting CAPON, the objective of this preliminary SAR was not extensive optimization, but rather confirmation of scaffold dependence and identification of essential pharmacophoric features. The limited number of derivatives evaluated reflects the early-stage nature of this study and the availability of close structural analogs within commercial libraries. This focused set was sufficient to establish structure-binding relationships and determine whether the **MA48** chemotype provides a viable foundation for future medicinal chemistry optimization. Analog selection was guided by systematic variation of three structural regions: (i) the thiazolothiadiazole core, (ii) the amino-based linker, and (iii) hydrophobic substituents on the terminal aromatic ring. These modifications were chosen to interrogate core scaffold integrity, hydrogen bond donor requirements, and hydrophobic pocket tolerance, respectively.

SAR analysis of the evaluated thiazolothiadiazole derivatives (**MA48, MA48.1, MA48.2, MA48.3, MA48.4, MA48.8, MA48.9, MA48.11**, and **MA48.13**) shown in (Table 1) revealed several key pharmacophoric features (Figure 3) that are crucial for the CAPON binding affinity. The pharmacophoric features consisted of a heteroaromatic core responsible for anchoring interactions, a hydrogen bond donor proximal to the core scaffold, and two spatially distinct hydrophobic regions that modulate affinity through aromatic and aliphatic interactions. Among all the tested derivatives, the thiazolothiadiazole motif was present in all active compounds and is therefore inferred to serve as the primary anchoring element within the binding site. The multiple ring nitrogens likely act as hydrogen bond acceptors or participate in electrostatic interactions, stabilizing the ligand-protein complex. Loss of CAPON binding affinity observed upon extensive substitution or rigidification of this core suggests a constrained binding pocket that favors minimal steric perturbation around the heterocycle seen in compound **MA48.13** (Kd > 200 μM, Table 1).

**Table 1.**
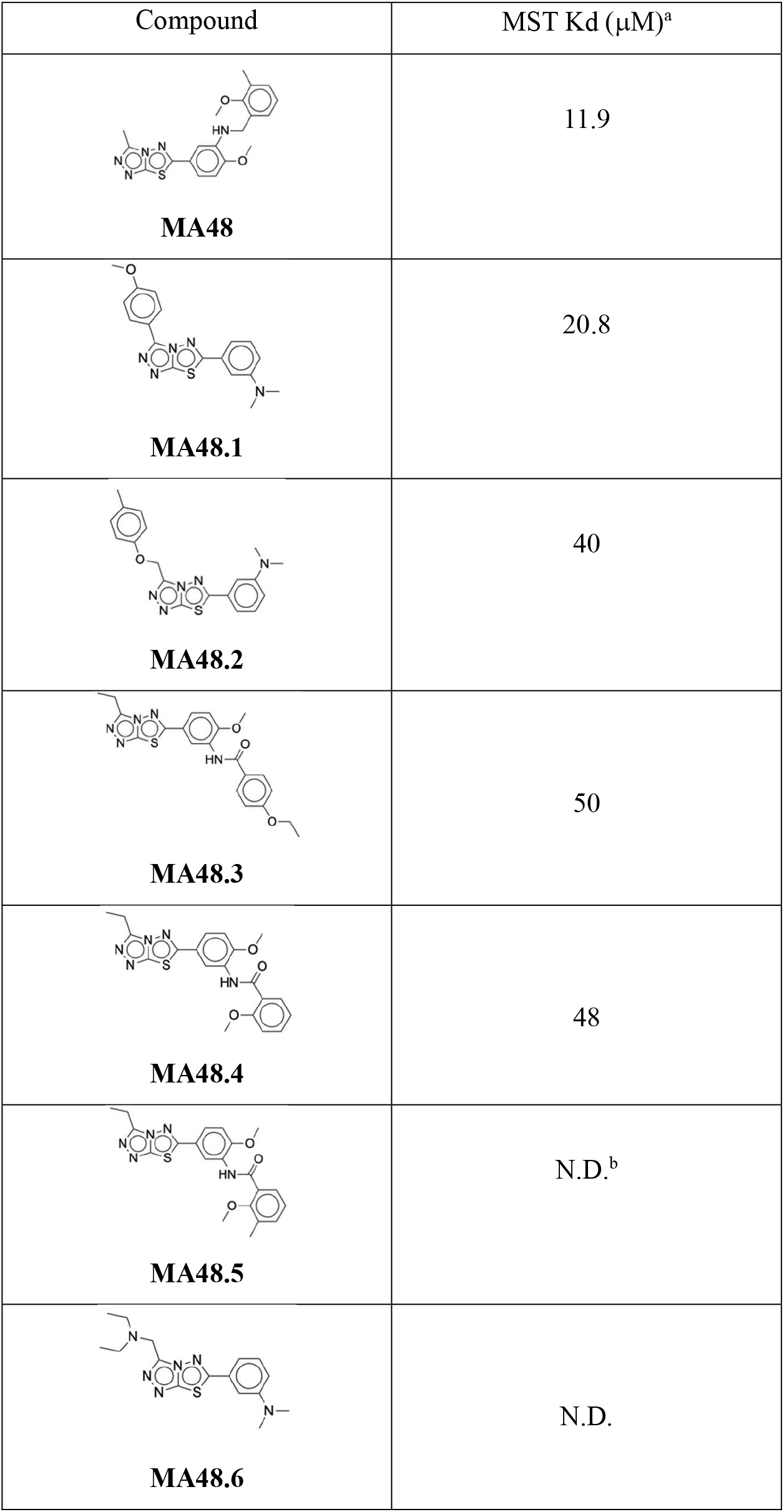
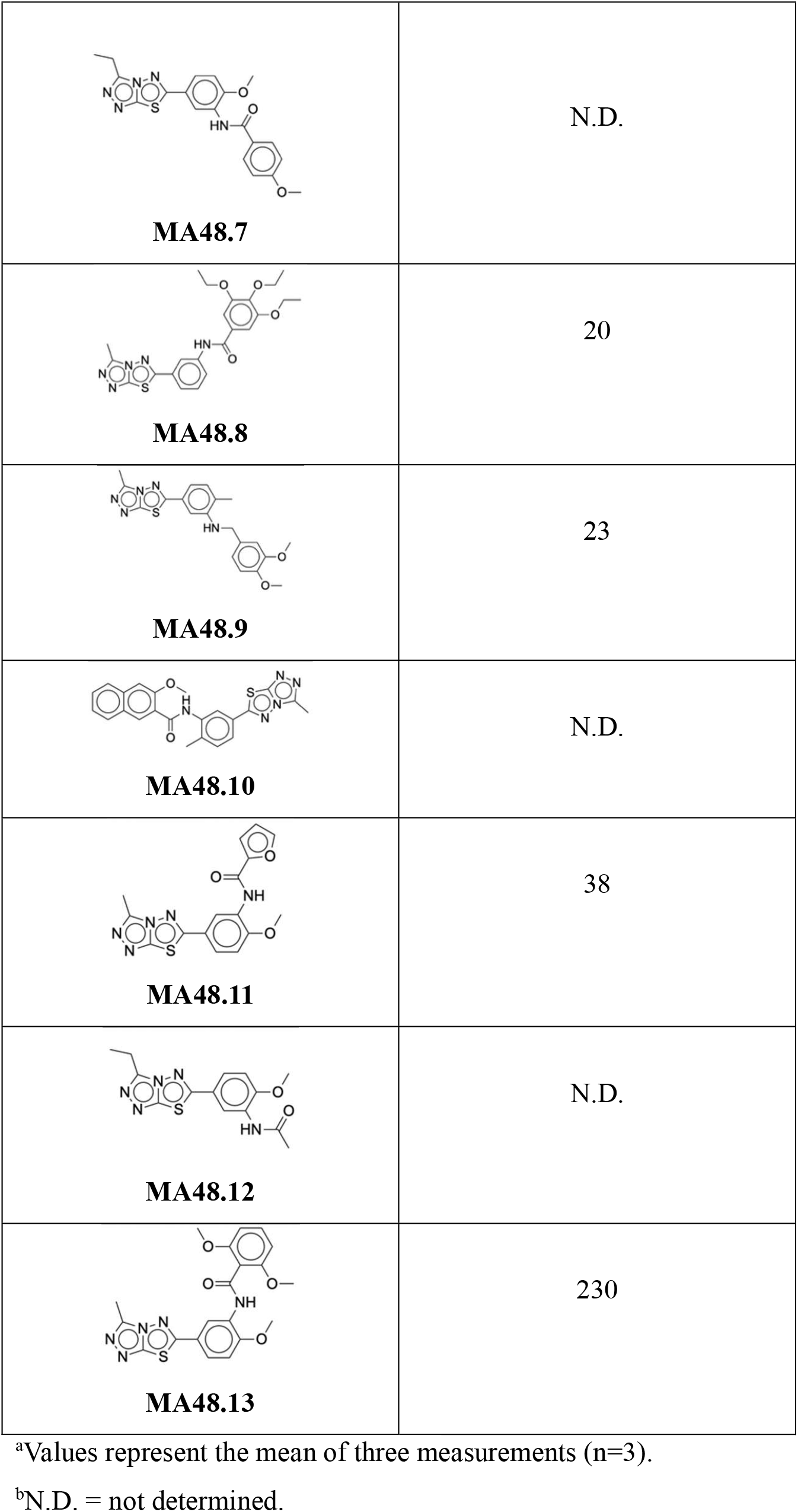
Chemical structures and CAPON binding affinities of compound **MA48** derivatives. Kd values were determined by microscale thermophoresis (MST).

| Compound | MST K<sub>d</sub> (μM)<sup>a</sup> |
| --- | --- |
| **MA48** | 11.9 |
| **MA48.1** | 20.8 |
| **MA48.2** | 40 |
| **MA48.3** | 50 |
| **MA48.4** | 48 |
| **MA48.5** | N.D.<sup>b</sup> |
| **MA48.6** | N.D. |
| **MA48.7** | N.D. |
| **MA48.8** | 20 |
| **MA48.9** | 23 |
| **MA48.10** | N.D. |
| **MA48.11** | 38 |
| **MA48.12** | N.D. |
| **MA48.13** | 230 |

<sup>a</sup>Values represent the mean of three measurements (n=3).
<sup>b</sup>N.D. = not determined.

**Figure 3.**
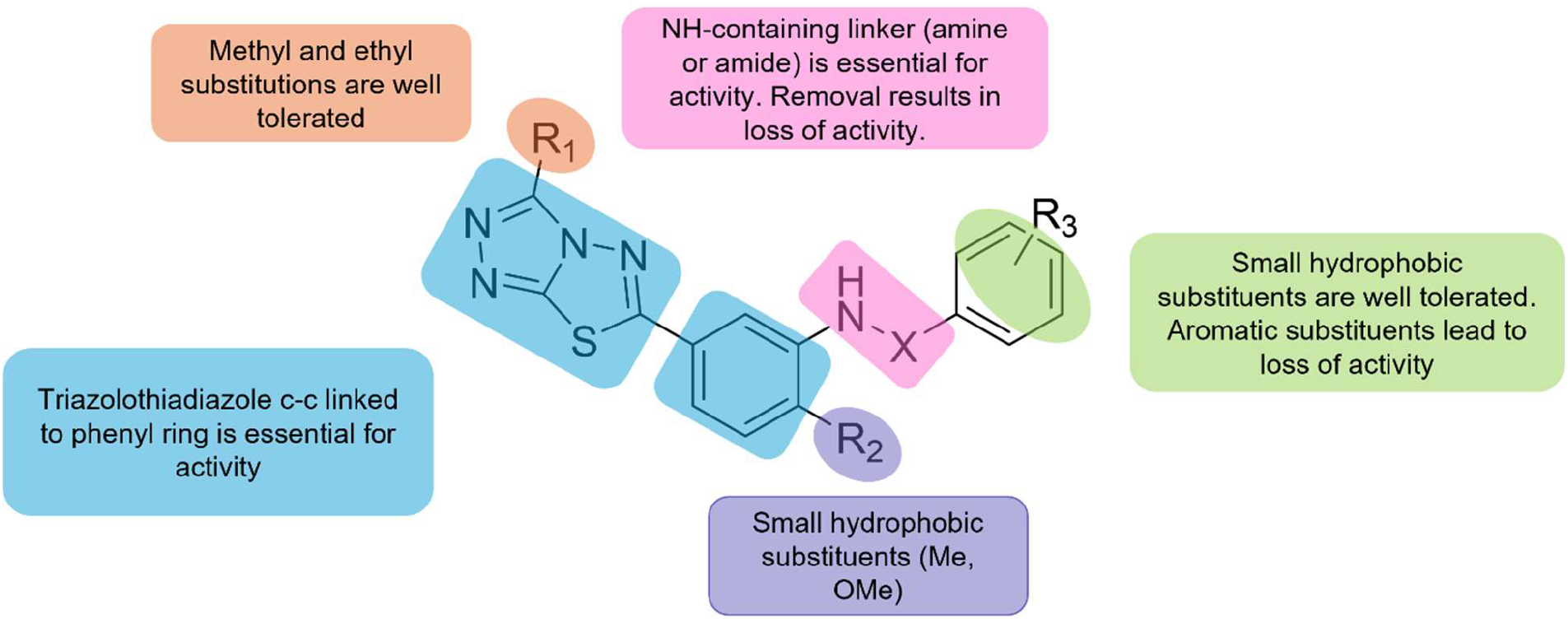
Preliminary structure-activity relationship (SAR) analysis of the MA48 chemotype. Structural modifications were introduced to probe the contribution of the thiazolothiadiazole core, the amino linker, and terminal hydrophobic substituents to CAPON binding affinity as determined by MST. Variations reveal the importance of the heterocyclic core and NH linker for maintaining measurable binding.

Another key pharmacophoric feature observed is the presence of an amino-containing linker (amine or amide-based linkers) connecting the core to a substituted aromatic ring. Loss or substitution of the amino-based linker results in markedly reduced binding affinity or complete loss of activity tested in derivatives **MA48.5, MA48.6, MA48.7, MA48.10** and **MA48.12** (Table 1). These observations suggest that this NH group forms a directional hydrogen bond with the target or that flexibility in this region is essential for the anchoring of the small molecules to the binding site. Hydrophobic interactions were similarly found to play a central role in affinity modulation. Two distinct hydrophobic regions were identified: a proximal aromatic moiety adjacent to the heterocycle and a distal hydrophobic substituent, typically a small alkyl or methoxy group as in **MA48**. Moderate substitution with small hydrophobic groups (e.g., methyl or methoxy) was well tolerated and, in some cases, enhanced binding, likely through favorable van der Waals interactions as depicted in derivatives **MA48.2, MA48.3, MA48.4** and **MA48.11**. In contrast, excessive aromatic bulk, increased rigidity, or highly substituted benzamides led to diminished activity, consistent with steric clash or entropic penalties within the binding site. Collectively, these data confirm that CAPON engagement is scaffold-dependent rather than nonspecific and establish the thiazolothiadiazole chemotype as a validated starting point for rational lead optimization.

Compounds bearing increased polarity or extended flexible side chains demonstrated intermediate activity (**MA48.2, MA48.3, MA48.4** and **MA48.11**), indicating that the binding pocket can accommodate limited polar functionality, provided that the core pharmacophoric features are preserved. However, excessive flexibility or over-polarization resulted in decreased affinity, underscoring the importance of maintaining a balance between conformational freedom and ligand preorganization. Collectively, these findings highlight the key pharmacophoric features that are key for maintaining the binding affinity. Future modifications that retain the NH linker while fine-tuning hydrophobic substituents are expected to yield compounds with improved potency and selectivity.

### Molecular Docking

Given that there are currently no crystal structures for CAPON protein, homology modelling was carried out to predict the structure of CAPON using the SWISS-MODEL server (Figure 4A). The platform identified the PTB domain-containing engulfment adapter protein 1 as the most suitable template, with 29.45% sequence identity. The GMQE score of 0.17 and QMEAND Global score of 0.57 ± 0.07 suggest moderate reliability of the predicted structure. The structural model reveals key features of CAPON, including helical and loop regions that may be important for protein-protein interactions or ligand binding. The predicted binding site identified by PrankWeb represents a potentially druggable pocket on the CAPON surface (Figure 4B). The successful docking of **MA48** into this predicted site (Figure 4C,D) suggests that this region may accommodate small molecules and could serve as a target for therapeutic intervention.

**Figure 4.**
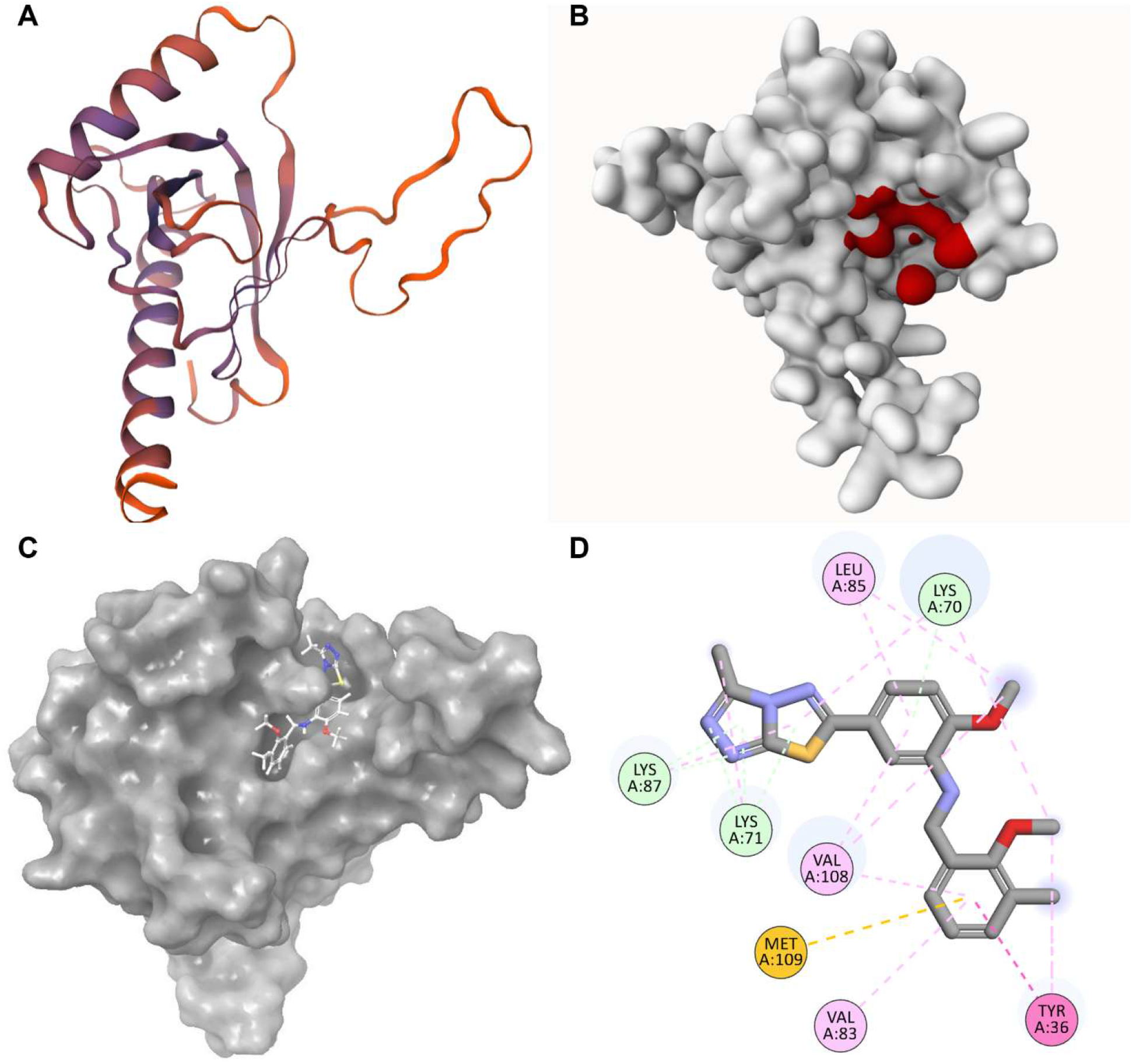
Structural Model and Ligand Docking Analysis of CAPON. **(A)** 3D representation of the CAPON homology model generated by SWISS-MODEL using the PTB domain-containing engulfment adapter protein 1 (PDB ID: 6itu.1.A) as template. **(B)** Surface representation of the CAPON model showing the predicted binding site (highlighted in red) identified using the PrankWeb server. **(C)** Surface representation of CAPON with the docked conformation of **MA48** positioned within the predicted binding pocket. **(D)** Two-dimensional interaction diagram of **MA48** bound to CAPON, generated using Discovery Studio Visualizer.

Analysis of the docking pose reveals several key interactions between **MA48** and residues in the CAPON binding pocket (Figure 4C,D). The interaction diagram shows contacts with multiple lysine residues (Lys70, Lys71, Lys87), leucine (Leu85), valine residues (Val83, Val108), methionine (Met109), and tyrosine (Tyr36). These interactions appear to involve a combination of hydrophobic contacts and potential hydrogen bonding, which together contribute to the predicted binding affinity. The presence of both charged and hydrophobic residues in the binding site suggests that the pocket can accommodate ligands with diverse chemical properties.

The central heterocyclic core of **MA48** appears to make favorable interactions with the binding pocket, while the aromatic substituents extend into hydrophobic regions. It is important to note several limitations of this computational study such as the moderate quality scores of the homology

model indicate that structural predictions. Another limitation to be considered is that the binding site prediction algorithm provides probable sites but does not guarantee that the identified pocket is functionally relevant or represents the true binding site for **MA48**. Despite these limitations, the computational pipeline employed here provides valuable structural insights that can guide experimental studies. The homology model offers a working hypothesis for the CAPON structure that can be refined as more experimental data become available. The docking results suggest that CAPON may be amenable to small molecule modulation and identify specific residues that could be targeted in mutagenesis studies or structure-activity relationship investigations. Notably, the observed SAR trends are consistent with the predicted docking pose, in which the heterocyclic core occupies a defined anchoring region while peripheral substituents extend into adjacent hydrophobic pockets.

### MA48 disrupts NOS1-NOS1AP interaction

To determine whether direct CAPON binding translates into functional modulation of the CAPON-NOS1 complex in a cellular context, we employed a NanoBRET-based protein-protein interaction assay in CHO-K1 cells co-expressing NanoLuc-tagged NOS1 and Venus-tagged NOS1AP. In this system, proximity between NOS1 and CAPON generates a measurable BRET signal, providing a quantitative readout of complex formation in living cells.

Treatment with **MA48** resulted in a concentration-dependent reduction in the NanoBRET signal (Figure 5), indicating disruption of the NOS1-CAPON interaction. At 100 μM, **MA48** reduced the NanoBRET ratio to approximately 80% of vehicle control, with statistically significant inhibition also observed at 50 μM. The cellular concentrations required for measurable disruption are consistent with the 11.9 µM biophysical affinity, taking into account factors such as intracellular exposure, protein abundance, and the dynamic nature of protein-protein interactions in live cells. Although the magnitude of disruption is modest, this degree of inhibition is consistent with the low-micromolar binding affinity measured by MST and is typical of early-stage scaffolds targeting protein-protein interfaces. Notably, complete dissociation of stable protein-protein complexes is not necessarily required to achieve biological modulation, and partial disruption is frequently observed in early-stage PPI-directed small molecules.

**Figure 5.**
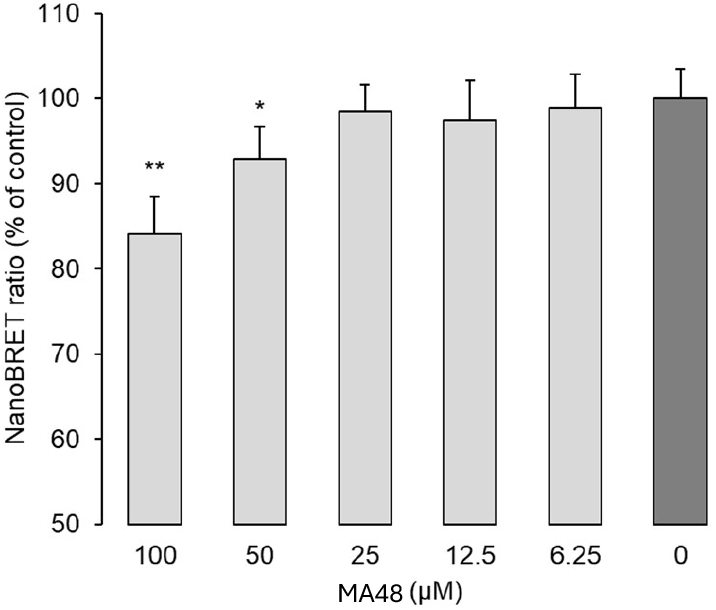
Compound MA48 inhibits NOS1-NOS1AP interaction in NanoBRET assay. CHO-K1 cells co-expressing NanoLuciferase-tagged NOS1 (N-terminal) and Venus-tagged CAPON (C-terminal) were treated with indicated concentrations of Compound **MA48**. NanoBRET ratio was measured and expressed as a percentage of the vehicle control. Data are presented as mean ± SEM (n = 5). *p < 0.05, **p < 0.01 vs. vehicle control (Student’s t-test).

Because NanoBRET measures proximity rather than direct binding, the observed signal reduction likely reflects decreased complex formation rather than destabilization of individual protein expression levels. Importantly, the observation that **MA48** perturbs the interaction in a live-cell context demonstrates that CAPON engagement by small molecules can translate into functional modulation of the NOS1-CAPON complex. These data provide proof-of-concept that the CAPON-NOS1 interface is chemically addressable and support further optimization of this chemotype to enhance potency and cellular efficacy. While broader selectivity profiling will be required in future studies, the concordance between biophysical binding and cellular interaction disruption supports a target-driven mechanism.

Adaptor proteins such as CAPON have traditionally been considered challenging targets for small molecule intervention due to their shallow interaction surfaces and lack of well-defined catalytic pockets. Demonstrating that the CAPON-NOS1 interface can be modulated by a drug-like small molecule therefore represents an important conceptual advance. Although **MA48** exhibits low-micromolar affinity consistent with early-stage PPI scaffolds, the identification of a tractable chemotype establishes CAPON as a chemically addressable target. This work lays the foundation for future medicinal chemistry optimization and for mechanistic studies aimed at dissecting CAPON-dependent signaling in neurodegenerative disease.

## ASSOCIATED CONTENT

### Supporting Information

The Supporting Information is available free of charge on the ACS Publications website.

Experimental data for ASMS (PDF)

CAPON homologue structure (PDB)

CAPON homologue data (PDF)

## AUTHOR INFORMATION

### Author Contributions

The manuscript was written through contributions of all authors. All authors have given approval to the final version of the manuscript.

### Notes

The authors declare no competing financial interests.

## Acknowledgments

This work was supported by the National Institute on Aging under grant number RF1AG084635(PI: Gabr).

---

Table of Contents artwork

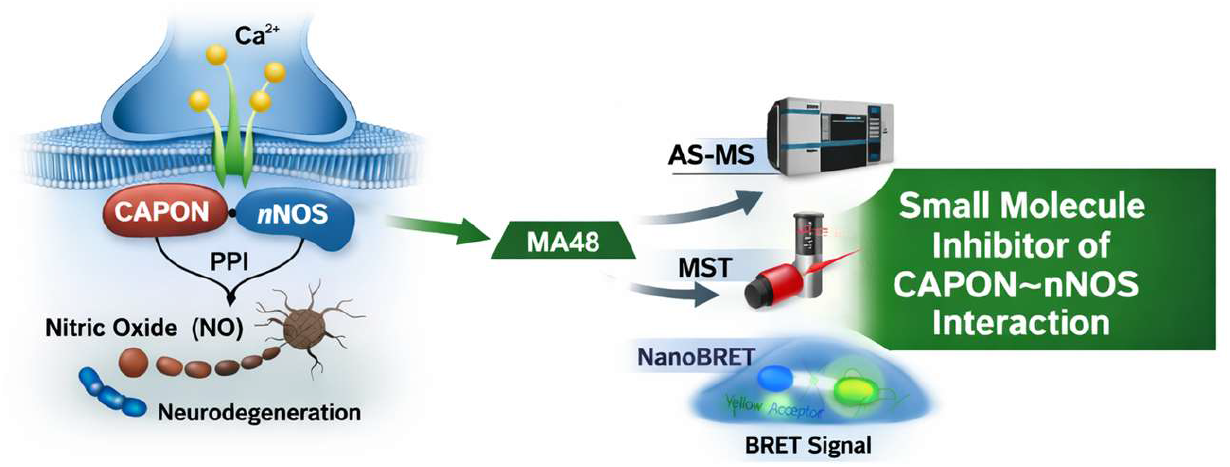

